# A Manifold-Based Framework for Studying the Dynamics of the Vaginal Microbiome

**DOI:** 10.1101/2023.09.06.556518

**Authors:** Mor Tsamir-Rimon, Elhanan Borenstein

## Abstract

The vaginal bacterial community plays a crucial role in preventing infections. The composition of this community can be classified into five main groups, termed community state types (CSTs). Four of these CSTs, which are primarily consisted of Lactobacillus species, are considered healthy, while the fifth, which is composed of non-Lactobacillus populations, is considered less protective. This latter CST is often considered to represent a state termed Bacterial vaginosis (BV) – a common disease condition associated with unpleasant symptoms and increased susceptibility to sexually transmitted diseases. However, the exact mechanisms underlying BV development are not yet fully understood, including specifically, the dynamics of the vaginal microbiome in BV, and the possible routes it may take from a healthy to a BV state. This study aims to identify the progression from healthy Lactobacillus-dominant populations to symptomatic BV by analyzing 8,026 vaginal samples and using a manifold-detection framework. This approach is inspired by single-cell analysis and aims to identify low-dimensional trajectories in the high-dimensional composition space. This framework further order samples along these trajectories and assign a score (pseudo-time) to each sample based on its proximity to the BV state. Our results reveal distinct routes of progression between healthy and BV state for each CST, with pseudo-time scores correlating with community diversity and quantifying the health state of each sample. BV indicators, including Nugent score, positive Amsel’s test, and several Amsel’s criteria, can also be successfully predicted based on pseudo-time scores. Additionally, Gardnerella vaginalis can be identified as a key taxon in BV development using this approach, with increased abundance in samples with high pseudo-time, indicating an unhealthier state across all BV-development routes on the manifold. Taken together, these findings demonstrate how manifold detection can be used to successfully characterizes the progression from healthy Lactobacillus-dominant populations to BV and to accurately quantify the health condition of new samples along the route of BV development.

## Introduction

The human body is inhabited by bacterial communities, collectively known as the human microbiome. These communities inhabit multiple sites, including the human gut, skin, and vagina. The composition of the vaginal bacterial population, specifically, can have a major impact on women’s health in multiple ways, for example, by interfering with the proliferation of harmful organisms^1,2^.

In an attempt to better understand and characterize the composition of the vaginal microbiome, researchers have sought to cluster the various compositions observed in this community into distinct groups. Such studies have shown that there are at least five major typical compositions of the vaginal microbiota, referred to as community state types (CSTs)^3,4^. Each CST is defined according to the dominant species in the community or the combination of species in this state. Specifically, several *Lactobacillus*-dominated compositions exist, including L. *crispatus-*, *L. gasseri-*, *L. iners-*, and *L. jensenii-*dominated communities, and are referred to as CST I, CST II, CST III, and CST V, respectively. Populations composed of other anaerobes (i.e., non-*Lactobacillus-*dominated), such as *Gardnerella*, *Prevotella*, and *Atopobium*, are referred to as CST IV. CSTs can be further separated into subgroups (subCSTs). For example, CST IV is further stratified into CST IV-A (dominated by *Lachnocurva vaginae*; BVAB), CST IV-B (dominated by *Gardnerella vaginalis*), or CST IV-C (a diverse collection of anaerobes)^5^. Similarly, within the *Lactobacillus*-dominant CSTs, CST I and III are divided into two subgroups, A and B. Subgroup A is composed of a higher abundance of the focal *Lactobacillus* species, while the subgroup B is composed of a lower abundance of that species. Finally, CST IV-C is also divided into 5 groups (labeled 0 to 4), when IV-C0 is a more diverse population and the other groups are dominated by a BV-related bacterium which is not *G. vaginalis* or BVAB.

Notably, different CSTs are also tightly associated with various physiological and clinical phenotypes. *Lactobacillus-*dominant CSTs, for example, are generally characterized by low levels of pro-inflammatory cytokines and low pH, which is ascribed to the production of lactic acid by the dominating *Lactobacillus* species^3,6^. Non-*Lactobacillus* CSTs, in contrast, can be accompanied by high pH and unpleasant symptoms, such as mal-odor and abnormal discharge. CST IV is often referred to as Bacterial Vaginosis (BV), a common dysbiotic condition. BV is also associated with multiple adverse gynecologic consequences, including an increased risk of preterm birth^7^, pelvic inflammatory disease^8^, acquisition of STIs including HIV^9^, and human papillomavirus^10^. Unfortunately, even though BV is very common, its etiology and development are still not clear, calling for a more principled mapping of its dynamics.

Indeed, the composition of the vaginal microbiome is not static, and changes throughout a woman’s life, from childhood to reproductive age, and during pregnancy and menopause^11^. These changes are ascribed to resource availability in the vaginal environment, such as the abundance of glycogen in the vaginal epithelium, which in turn seems to be highly affected by estrogen levels^12^. Changes may also occur at much shorter timescales, for example, in correspondence to the menstrual cycle^13^. Sharp compositional shifts can also be observed, for example, following sexual intercourse and other host habits^13–16^. These various processes and factors are clearly not independent, making the dynamics of the vaginal microbiome challenging to comprehensively characterize, in spite of the relatively simple composition of that microbiome.

Clearly, while changes in the composition of the microbiome occur in a high-dimensional space (i.e., spanning all possible community compositions), microbial dynamical processes (such as the transition from a healthy state to BV) may potentially follow relatively common patterns, and accordingly, likely progress along a low-dimensional trajectory embedded in this high-dimensional space. The detection and characterization of such trajectories, however, is a challenging task as the vaginal microbiome is affected and may be perturbed by many factors. Indeed, several approaches were used to describe the dynamics of the vaginal microbiome. Gajer *et al.*, for example, have used an ordination analysis to show a low dimension representation of vaginal microbiome compositions (based on cross-sectional and longitudinal data) and the progression of each woman’s microbiome over time^3,17^. This study clearly illustrated the continuous nature of transitions between CSTs, albeit, without a qualitative interpretation of intermediate stages. Other studies used integration of time-series data across individuals by alignment of longitudinal vaginal samples^18,19^. The aligned data in these studies revealed some interesting insights, including, for example, an antagonistic behavior between *L. iners* and *Atopobium*. An additional longitudinal metagenomic study of healthy women with high risk for BV proposed that BV formation is initiated by early colonizers (such as *Gardnerella vaginalis*), creating a more favorable environment for other BV-related bacteria^20^. However, to date, a rigorous approach for describing the route of transition between *Lactobacillus*– and non-*Lactobacillus-* dominant CSTs has not yet been presented, calling for a robust framework that enables identification of complex dynamical processes in this high-dimensional data.

Interestingly, dynamical processes and progression trajectories can be potentially identified not only via longitudinal data analysis, but also via the analysis of massive cross-sectional datasets. Such datasets can be viewed as a collection of snapshots into the underlying dynamical process, such that each sample provides information about a specific point along the progression trajectory. Identifying low-dimensional trajectories embedded in this high-dimensional space and ordering samples along the identified trajectories can therefore both help characterizing the dynamical processes involved and enable labeling samples according to their location along such processes. An example of this approach has been presented in a study by Li *et al.*, who suggested a model that recapitulated longitudinal progression of the gut microbiome in Crohn’s disease^21^. Based on a combination of clustered samples and a principal tree (which represents the topology of trajectories and alterations in the microbial composition), they proposed a double bifurcating model of microbial alterations that occur during Crohn’s development. While this model successfully captures the complex structure of disease progression, the low sample size limits the model’s robustness and applicability. Another study applied the pseudo-time approach to examine the structure of the gut microbiome throughout the human lifespan^22^. This research discovered distinct clusters within the microbial structure that were linked to the proportions of specific bacterial genera, along with associations between several functional characteristics and the endpoints of these clusters. While yielding intriguing findings, the pseudo-time analysis did not uncover insights into the progression of diseases, potentially owing to the complex composition and high variation of the gut microbiome. Similar analyses of dynamical processes in complex biological systems are also common in single-cell studies, where high dimensionality and high resolution data provide snapshots of single cell states along some processes, such as cell maturation or differentiation^23,24^. Such analyses then aim to identify the low dimensional trajectory (or ‘manifold’) embedded in the high dimensional space of cell states and label each cell according to its location along this trajectory.

In this study, we similarly use a framework inspired by single-cell analysis approaches for manifold detection to identify and characterize the dynamics of the vaginal microbiome. Specifically, we aim to characterize the trajectories of progression from *Lactobacillus*-dominant populations to non-*Lactobacillus* population. Identifying low-dimensional trajectories in the vaginal microbiome composition space and projecting new cross-sectional or longitudinal samples on these trajectories allow us to both map the potential routes of the vaginal microbiome and to trace the dynamic along these trajectories (e.g., by examining the progression of a certain woman along these trajectories over time). Moreover, combining cross-sectional and longitudinal data allows us to quantitatively characterize the dynamics of various women, and to compare, for example, the progression along various trajectories to multiple clinical variables. Combined, our findings provide a more principled perspective for examining the vaginal microbiome dynamics, including complex transitions between CSTs and BV development.

## Results

### A manifold detection approach for characterizing the dynamics of the vaginal microbiome

Aiming to comprehensively characterize the dynamics of BV development, we utilized a computational framework that both identify various possible low-dimensional routes in the microbiome compositional space, and place each sample along these routes. This framework first takes as input a large collection of microbiome samples (from either cross-sectional or longitudinal studies; Fig. 1a) and utilizes a previously introduced manifold detection algorithm^25^ to reduce the dimensionality of the data and to identify a low-dimensional manifold that describes the distribution of all input samples in this space (Fig. 1b). This algorithm further labels each sample according to its location on the manifold and the distance of this location to a pre-defined set of root samples (Fig. 1c). This label, referred to in the single-cell literature as pseudo-time, quantitatively assess how close, along the manifold’s available routes, is the sample of interest from some given state. In our analysis, we used highly diverse BV samples (mostly assigned with CST IV-B) as root and defined pseudo-time between 0 and 1, such that 0 denotes a sample far from the BV state (the “healthiest” composition) and 1 as the root (a BV composition). Specifically, in this study, we used the PAGA (Partition-based graph abstraction) manifold detection algorithm^25^, which projects the samples on a low dimensional space and then utilizes the distance matrix based on the low dimensional graph to produce pseudo-time labels (see Methods for full details).

**Figure 1:**
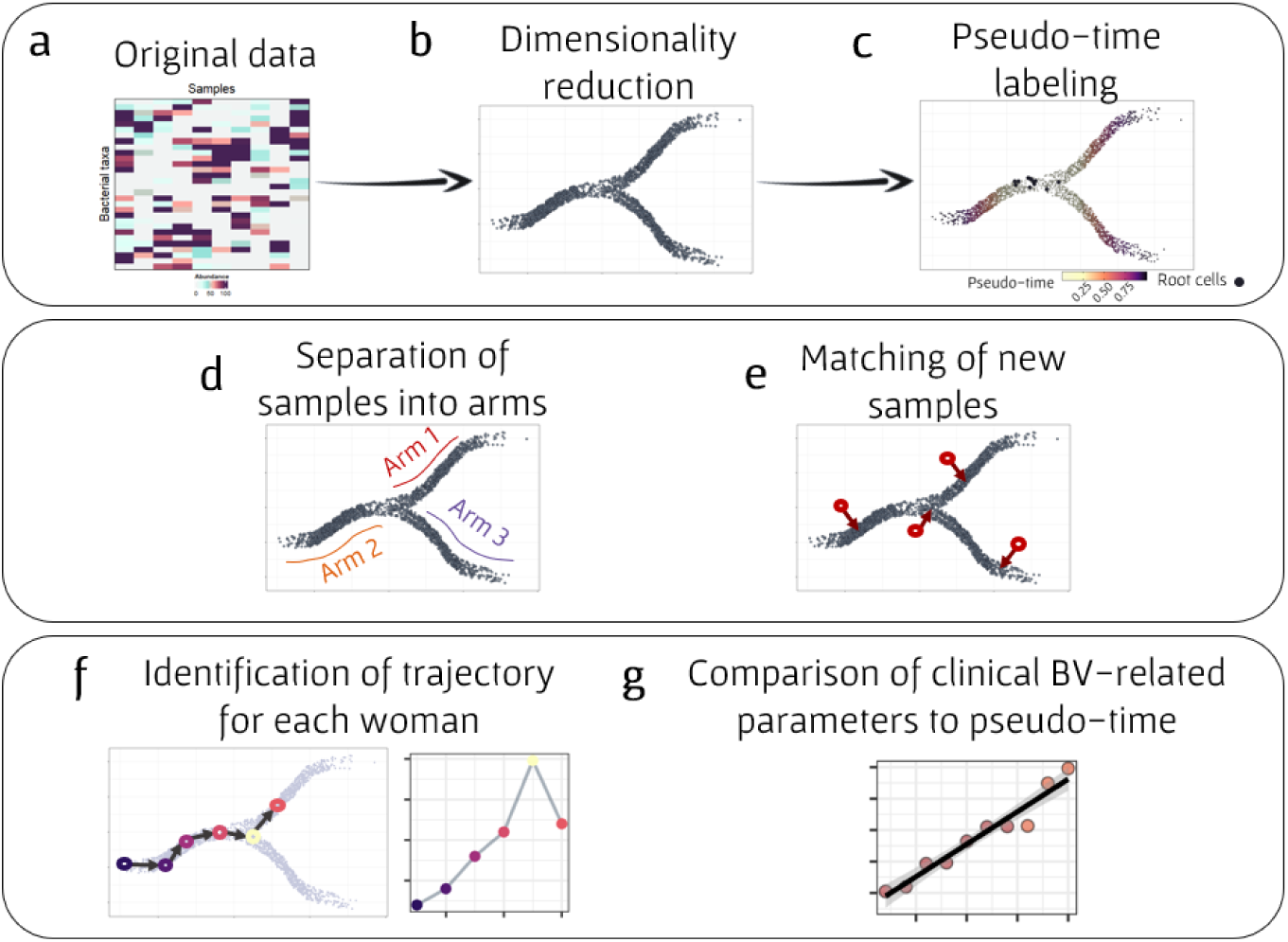
Analysis scheme of the manifold detection framework. **(a)** The framework takes as input a high dimensional microbial data table, describing microbial features of each sample. **(b)** Data is then projected onto a lower dimensional space, resulting in a compositional manifold. **(c)** Each sample is labeled with a pseudo-time score, calculated based on its distance along the manifold from a pre-defined group of root samples. **(d)** The manifold is also partitioned into “*arms*” (which represent distinct routes on the manifold and different possible routes to BV), based on the assigning CSTs to the various sample. **(e)** New samples are matched to their nearest neighbor on the manifold, obtaining their pseudo-time label and location on the manifold from their nearest neighbor. **(f)** Samples from a longitudinal dataset can be mapped to this manifold and used to characterize the trajectory of each woman along the manifold, comparing samples’ pseudo-time labels to the chronological time in which the sample was obtained. **(G)** Pseudo-time labels associations with various clinical BV-related variables, including community diversity, Nugent score, and pH, can be examined.

In addition, we also use a previously developed method for determining the CST of each sample^5^. Since samples from a given CST are also generally more similar to one another than to samples from a different CST, sample from each CST tend to form a distinct route (or an “arm”) in the above identified manifold (Fig. 1d). Finally, given a new set of samples (i.e., that were not part of the input and the detected manifold, such as samples from some women in a new longitudinal study), we match each new sample to its nearest neighbor sample on the manifold, using the location and pseudo-time label of this nearest neighbor to label the new sample (Fig. 1e).

Combined, this procedure assigns each sample a quantitative measure of its progression along possible routes from healthy to BV states, allowing us to study the dynamics of the vaginal microbiome. For example, focusing on longitudinal samples from a single woman, and comparing the pseudo-time label of these samples to the chronological time in which they were obtained, allow us to comprehensively map each woman’s trajectory along the route of BV development (Fig. 1f). In addition, we can compare samples’ pseudo-time labels to various clinical parameters, to examine correspondence with BV progression (Fig. 1g), or use pseudo-time to predict clinical BV-related variables.

### Analysis of vaginal community dynamics in a large-scale dataset collection via manifold detection framework

To assess the ability of the framework described above to identify BV developmental trajectories in the vaginal microbiome, we analyzed five different datasets, including three cross-sectional datasets and two longitudinal datasets. Briefly, the cross-sectional datasets, combined, include thousands of samples, mostly from reproductive-age asymptomatic women. The longitudinal datasets, include again thousands of samples combined, with one dataset of 83 women sampled daily (with ∼70 samples per woman), and another of 84 women sampled bi-monthly (with 2-7 samples per woman). Sampled with insufficient coverage were filtered and discarded from downstream analyses. In total, our dataset collection after filtration included 8,541 samples. Available metadata varied across datasets, with some datasets providing detailed data about age, ethnicity, and various clinical BV indicators and criteria, while others included more limited metadata. Complete information about these datasets and available pertaining information can be found in the Methods.

The CST of each sample in our dataset collection was determined using the VALENCIA algorithm^5^. Across all five datasets, 57% of the samples were assigned to *Lactobacillus*-dominant CST, in general agreement with other studies^3,26^ (Supplementary Fig. S1a). CST assignment also produced a similarity score for each sample, indicating how much the sample resembled the core CST as defined by VALENCIA^5^ (see Methods). The mean similarity score across all samples was 0.75, and varied between CSTs, with mean similarity of 0.86 in the main *Lactobacillus* CSTs (I-A, II, III-A and V; Supplementary Fig S1a), 0.79 in the secondary *Lactobacillus* CSTs (I-B and III-B), and 0.56 in non-*Lactobacillus* CSTs (IV-A, IV-B and IV-C), again in accordance with similarity scores reported by other studies^27^. We also calculated the Shannon diversity index for each sample. As expected, Shannon diversity index varied substantially between communities, with CST IV-C0 and IV-A exhibiting the highest mean diversity (2.75 and 2.24, respectively), and CST I-A and III-A the lowest diversity (0.29 and 0.47, respectively; Supplementary Fig. S1b). These values match variation in diversity across CSTs as previously reported^5^.

Additionally, to allow us to evaluate the capacity of our framework to assign meaningful pseudo-time labels to “new” samples, we held out a set of samples from our dataset, using only the remaining samples for identifying and characterizing the manifold as described above. Specifically, we randomly selected 30 women from one of the cross sectional datasets (AVPVC)^3^ and 12 women from one of the longitudinal datasets (UAB^28^; each with multiple daily samples), for a total of 515 samples. These *held-out* samples were not included in our pseudo-time analysis, and instead were each mapped to its nearest neighbor (using the Bray Curtis distance metric) on the manifold and assigned the pseudo-time and CST of that nearest neighbor.

Having constructed and processed this dataset collection, we finally applied the framework described above to samples from the five datasets. Dimensionality reduction, as a part of the PAGA package, resulted in a graph with distinct “arms”, each could be assigned to a specific CST (Fig. 2a, inset). The arms of CST-I and III were the longest, and each can be further partitioned into the main subCST, A (with sample located closer to the periphery of the graph), and the secondary subCST, B (with samples closer to the center of the graph and to BV state samples). This illustrates the gradual changes in vaginal community along routes of BV development, as well as the prevalence of these CSTs^3,5^. Notably, samples from the five different datasets were relatively well-spread across the UMAP space (Fig. 2b), confirming that the result manifold is not an artifact of some study-specific bias. This manifold was then used to calculate the pseudo-time of each sample, based on provided root samples (Fig. 2c; see Methods). In our analysis, we defined samples with positive Amsel’s test (i.e., three positive criteria out of four) and with Shannon diversity index >3.5 (considered less healthy), as root samples.

**Figure 2:**
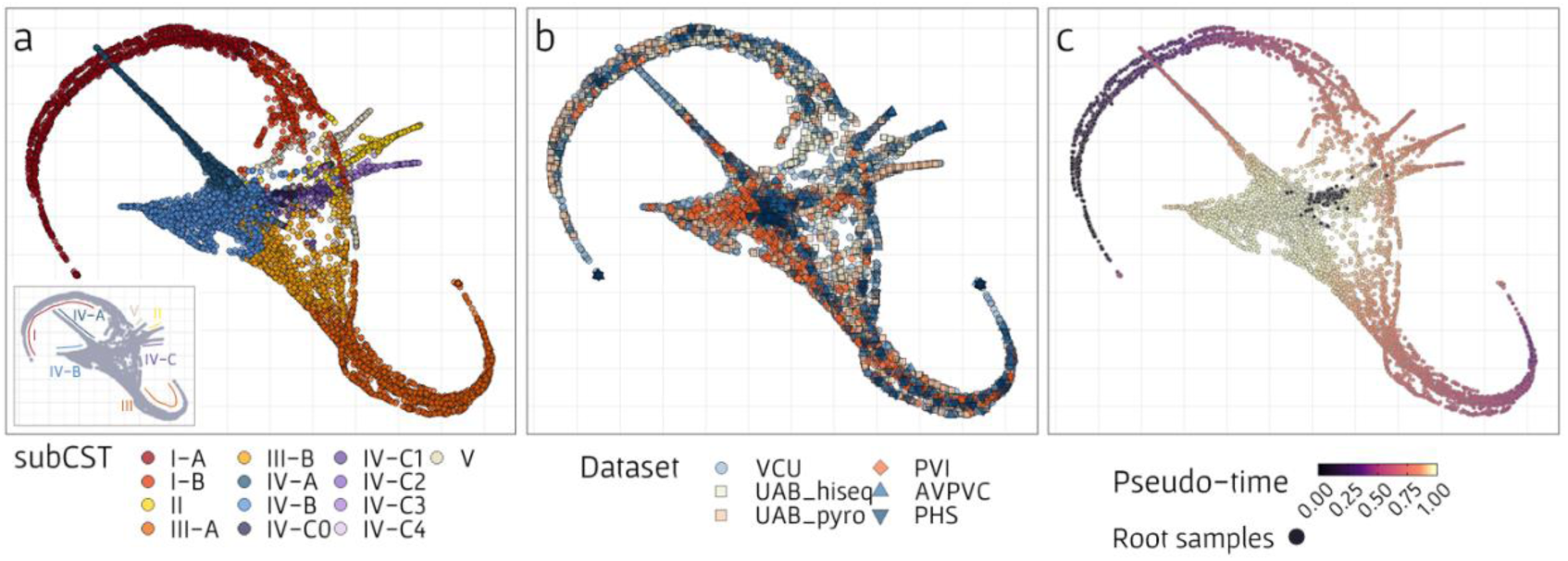
Visualization of the vaginal microbiome composition manifold and its relation to various sample properties. Each point represents a single sample (from the pooled set of the five datasets analyzed), where its location was determined by low dimensionality reduction using the PAGA algorithm. **(a)** Colors indicate the subCST determined for each sample based on VALENCIA. The inset on the bottom left corner represents the partition of the samples into the different manifold’s arms, where each arm is associated with a different CST. **(b)** Colors indicate the dataset each sample was obtained from. The UAB dataset was further partitioned according to the sequencing method that was used. **(c)** Colors indicate the pseudo-time label that was assigned to each sample based on results from the PAGA algorithm. Black points denote root samples, defined as highly diverse samples (Shannon diversity index >3.5) with positive Amsel’s test.

### Pseudo-time labels’ association with BV-related clinical indicators

One of the potential advantages and application of our pseudo-time approach is that it provides an objective quantification of a sample’s progression towards a BV state, based solely on observed trajectories along the vaginal microbiome compositional space. Indeed, for example, we found that the mean pseudo-time of *Lactobacillus*-dominant CSTs was 0.72, whereas that of non-*Lactobacillus* CSTs was 0.91, indicating that *Lactobacillus* populations are further away from an unhealthy state, as expected. To further confirm the utility of the obtained pseudo-time label as markers for BV progression, we next examined how they correlate with various ecological and BV-related features, such as Shannon diversity index, Nugent score, and Amsel’s criteria. We first focused on community diversity, acknowledging that a diverse vaginal population is generally considered less healthy and closer to BV state. This analysis was conducted separately for each arm on the manifold (based on Fig. 2a), since BV developmental patterns may vary across different source CSTs. As expected, we found that Shannon-diversity index was highly correlated with pseudo-time across all arms, with R^2^ of 0.79, 0.66, 0.8 and 0.69 in arms of CST I, II, III and V, respectively (Fig. 3a). Moreover, held out samples (see above) that were assigned with pseudo-time labels according to their nearest neighbors’ labels have also exhibited high correlation between their Shannon diversity values and the assigned pseudo-time label in the healthy available arms (CST I and III; Supplementary Fig. S2). Since root samples in our manifold were also defined as those with the highest Shannon-diversity index (along with positive Amsel’s test result), and other CSTs are generally defined by the dominance of one phylotype (and are hence less diverse), these correlations are not necessarily surprising, yet, the strong correlation suggests a good fit of the samples’ placement along the manifold and their bacterial population’s composition. Importantly, however, examining the relationship between pseudo-time and clinical BV indicators, we again found similar, albeit somewhat weaker correlations. Specifically, Nugent score in all arms, except for IV-C, exhibited a significant correlation with pseudo-time values (Spearman correlation; FDR <0.05). Furthermore, pseudo-time labels significantly differed between samples with positive and negative Amsel’s test results in arms I, III, IV-A, and IV-B. Arms II and V were excluded from this analysis due to insufficient sample size, while arm IV-C showed no significant difference (Supplementary Table 1). As for Amsel’s specific criteria, we found that pseudo-time showed significant correlations with pH in three arms (III, IV-B, and IV-C) and significant differences between positive and negative clue cells, whiff, and abnormal vaginal fluid tests in arm III.

**Figure 3.**
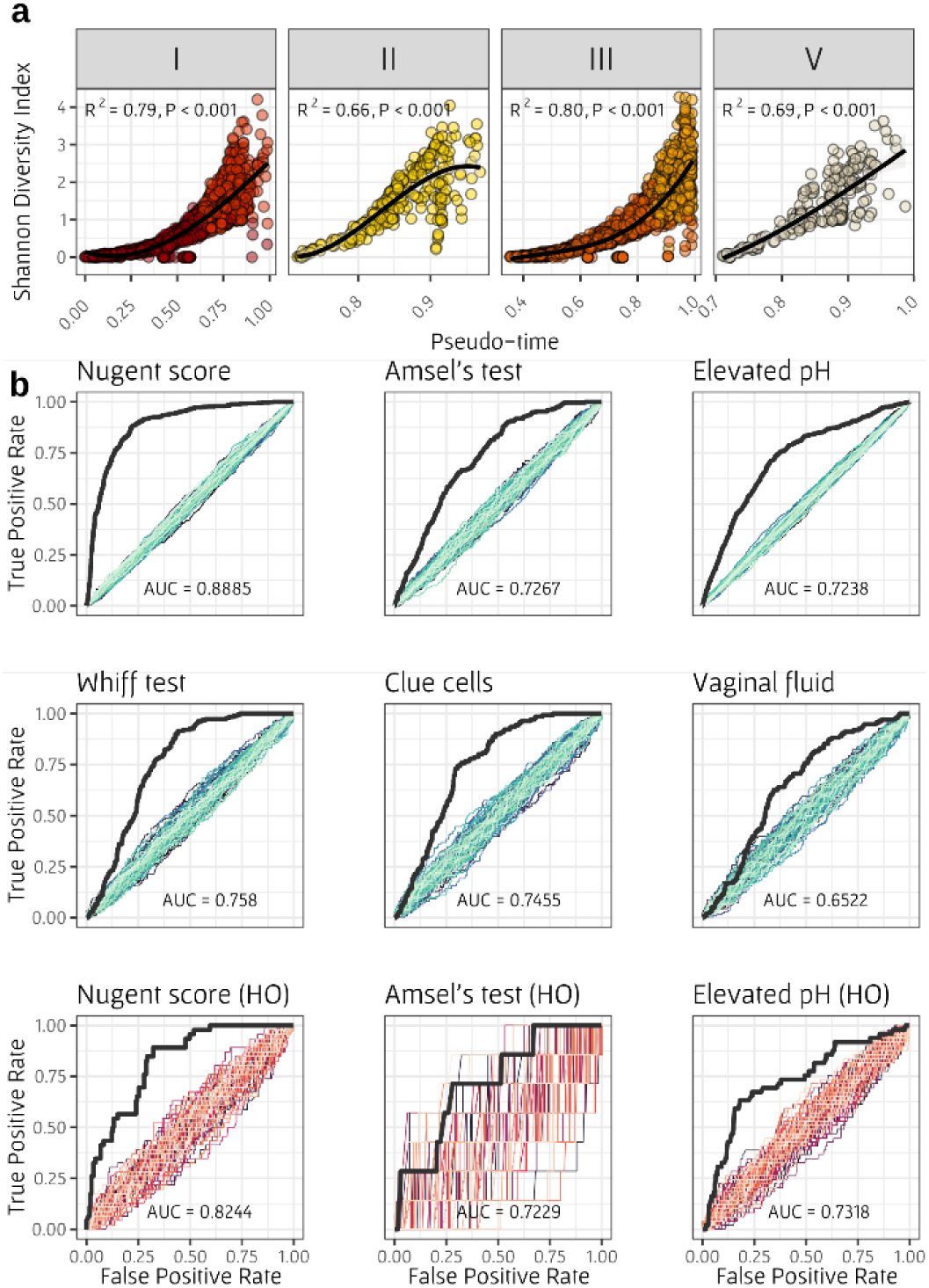
(a) Shannon diversity index as a function of pseudo-time label in healthy arms. Each plot represents a different CST arm and colors indicate subCST. The black line represents a linear regression to transformed polynomial values of pseudo-time. The R^2^ and p-value presented in each panel were determined by Spearman correlation test. **(b) Receiver operating characteristic (ROC) curves for predicting six BV indicators with pseudo-time labels**. The area under the ROC curve (AUC) is presented at the bottom of each plot. Colored curves represent prediction obtained using shuffled indicator labels. Bottom plots represent similar analysis, based only on held-out samples.

Given our primary objective, which is to elucidate the development of BV through pseudo-time analysis, we next aimed to examine how well BV-indicators can be predicted using pseudo-time labels. To this end, we calculated for each BV-indicator a Receiver Operating Characteristic (ROC) curve, describing how well can the pseudo-time label separate between positive and negative values of this indicator given different pseudo-time thresholds (Fig. 3b). Then, the area under the ROC curve (AUC) was calculated to quantify the overall predictive accuracy. To further assess the information captured in the pseudo-time labels, we compared the obtained ROC AUC to predictions using shuffled labels. Notably, samples defined as root samples for the manifold detection analysis were discarded from this analysis. Here, we focused on six BV indicators: high Nugent score, positive Amsel’s test, and the four criteria of Amsel (elevated pH, positive whiff test, presence of clue cells, and abnormal vaginal fluid). Since not all indicators were available in all datasets, we limited the analysis of each indicator to the dataset in which it was provided. Specifically, Nugent score was available only in the UAB study, Amsel’s test and pH were available in the PHS, UAB, and PVI studies, and the remaining Amsel’s criteria were available only in the PHS and PVI datasets. Evidently, our analysis revealed that pseudo-time could effectively predict all BV indicators, with ROC AUC ranging between 0.65 (and for all indicators except vaginal fluid, 0.72) to 0.88. The relatively low predictability of vaginal fluid, may be attributed to the dispersed distribution of samples with abnormal fluid on the manifold, and in agreement with one dataset from the original PVI study, where only BVAB1 was associated with abnormal vaginal fluid, compared to more than ten taxa significantly associated with other Amsel’s criteria^29^. Repeating this analysis using only the held-out samples, we observed similar predictability (since held-out samples were chosen from the AVPVC and UAB datasets, only the high Nugent score, positive Amsel’s test, and elevated pH indicators were available, with only 7 samples positive for Amsel’s test).

### Pseudo-time labels’ association with menstrual cycle and BV-related metabolites

Given the significant influence of the menstrual cycle on fluctuations in vaginal population composition^30,31^, we next aimed to investigate the relationship between pseudo-time labels and menstruation. We hypothesized that pseudo-time would exhibit greater fluctuation during the menstruation and period close to menstruation in comparison to fluctuations in other times. To investigate this, we calculated the difference in pseudo-time between each two consecutive samples, and compared the observed differences between menstruation (which we defined as the period ranging from two days before menstruation to two days after menstruation) and all other times. Our analysis indeed demonstrated a statistically significant difference in fluctuation between mensuration and non-menstruation periods, among the samples used in our manifold detection analysis (Wilcoxon test, Supplementary Fig. S5). A similar trend was observed in the held-out samples, but did not reach statistical significance.

In order to validate the effectiveness of our pseudo-time framework, we next examined the relationship between pseudo-time labels and BV-associated metabolites, such as biogenic amines^32–35^. These amines include, for example, the metabolites cadaverine, putrescine, and tyramine, which were previously linked to increased likelihood of transitioning from *Lactobacillus*-dominant CST to non-*Lactobacillus* CST. Indeed, data about the level of five biogenic amines (cadaverine, putrescine, spermine, spermidine, and tyramine), were available for samples from the UAB dataset used in our pseudo-time analysis^33^. We found the levels of cadaverine, putrescine, and tyramine (but not spermidine and spermine) were significantly higher in samples labeled with high pseudo-time (>0.9), compared to samples with low pseudo-time (P = 0.00001, 0.00029, 0.0044, respectively; Wilcoxon test; Supplementary Fig. S6), in agreement with the association between BV and these three biogenic amines reported above^32–35^. Combined, these analyses reveal robust correlations between pseudo-time labels and multiple key indicators of BV and BV-related metabolites.

### Individual trajectories of women from the longitudinal dataset along the manifold

Since our dataset collection includes longitudinal samples, wherein multiple samples were obtained from some women over time, we compared for each such woman the calculated samples’ pseudo-times with the chronological time at which the samples were obtained, thereby characterizing the specific trajectory of each woman along the manifold. Importantly, our pseudo-time calculation did not utilize information about which sample originated from which woman or at what chronological time in any way. As evident from top plots in figure 4, observed trajectories vary substantially between women. For example, woman UAB103 (Fig. 4a) demonstrates a relatively stable trajectory, remaining on the same healthy arm of the manifold (CST I) throughout the experiment’s duration, although moving back and forth (i.e., closer and further from the BV state) along this arm. In contrast, woman UAB059 (Fig. 4b) exhibits recurrent transition from a healthy state to a BV state (generally in sync with the menstrual cycle), moving from one arm to another. Finally, woman UAB002 (Fig. 4c) demonstrates frequent sharp transitions, moving between multiple arms on the manifold (including three different healthy arms and two BV-associated arms) without any discernible pattern. The trajectories of all women from this dataset on the manifold can be found in Supplementary Fig. S3, and pseudo-time progression along chronological time can be found in Supplementary Fig. S4.

**Figure 4:**
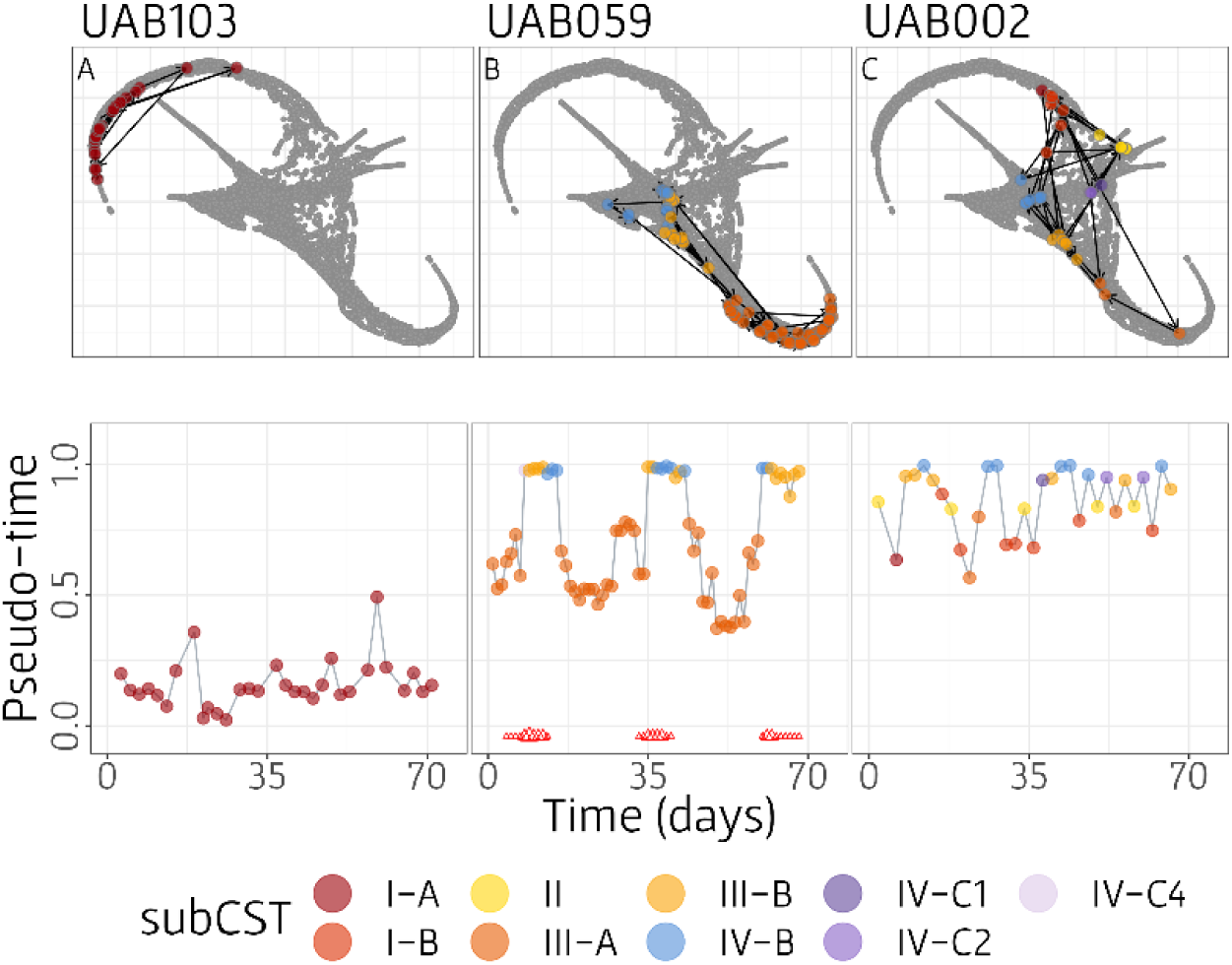
A manifold-based characterization of the trajectories of 3 representative women from the daily longitudinal dataset (UAB). The top plot in each panel illustrates the trajectory of the woman on the detected manifold. Grey dots represent the UMAP visualization of the vaginal microbiome composition manifold, and colored dots represent the locations of samples obtained from a specific woman, where each color represent the determined subCST of the sample. Black arrows denote the direction of the woman’s trajectory, showing the order of the samples based on the time they were obtained in the experiment. The bottom plot in each panel illustrates pseudo-time progression of the woman, as a function of chronological time. Dots colors are as in the top plots. Red triangles at the bottom of the plot represent self-reported menstruation, where size indicates menstruation spotting (small), medium or heavy bleeding (large).

These observed differences in women’s trajectories may reflect various factors affect community behavior. For example, a continuous increment in pseudo-time along chronological time (as seen, for example, in subjects UAB001; Supplementary Fig. S4) may represent a gradual progression towards BV, whereas, a sharp increase in pseudo-time (as seen in subject UAB075) might imply a specific event or exogenous perturbation that caused this shift. One such perturbation, for example, may be the use of medications for treating BV, aiming to shift the microbial population towards a healthier state. Although the efficacy of such medications can vary among women^36,37^, we hypothesized that the use of such medications should lead to a decrease in pseudo-time (i.e., a movement along the manifold towards a healthier state). To test this hypothesis and investigate the impact of BV medication on pseudo-time, we used available data from the UAB dataset, and analyzed samples collected during medication use, as well as one day before and after treatment. Among the ten instances of BV medication usage across 13 incidents of 11 women, nine indeed showed a consistent decrease in pseudo-time during this period (Supplementary Fig. S7), highlighting the potential of our pseudo-time labels as an effective model of temporal microbiome dynamics.

### Shifts in taxa abundances during BV development

Even though BV is a highly common disease worldwide, its etiologic agent is yet unknown. Specifically, our understanding of disease development in women with recurrent BV or with slow progression towards disease (i.e., in women not affected by sexual intercourse or another unique event^38^), is lacking. The vaginal microbiome composition manifold, presented in Figure 2, can serve as a comprehensive map of the potential routes of the vaginal microbiome, allowing us to quantify changes in bacterial abundances along various manifold’s arms, each representing a different route of BV development. For this purpose, samples were separated again into the different arms and ordered by pseudo-time label. To address the noisy and stochastic nature of the data, we further used a sliding window and calculated the average relative abundance of each taxon along pseudo time (see Methods; Fig. 6). An increase in a taxon’s abundance with pseudo-time values implies a potentially important role in BV development, especially if this increase is observed in multiple arms. For example, *G. vaginalis, Finegoldia,* and *Streptococcus* increased in abundance with pseudo-time in all arms. *G. vaginalis,* specifically, increased in abundance substantially with pseudo-time, reaching relative abundance of 2.3-11% in the samples with the highest pseudo-time, although these samples are still classified as being in a healthy CST. This finding supports the association between *G. vaginalis* and BV that was previously shown^43^, and suggests it has an important role in BV development from all healthy CSTs. Interestingly, all *Lactobacillus* species also tend to increase in abundance in the arms in which they were not the dominant species, potentially owing to a general increase in diversity and a corresponding shift from a single *Lactobacillus* species-dominant community to a more diverse community. Lastly, the abundance of several taxa increased in specific arms, such as *Corynobacterium* (arms I, II and V), *Bifidobacterium* (II and V), *Anaerococcus* (I and II), *Prevotella* (II), and *Megasphaera* (III). Even though these taxa’s abundances were relatively low throughout the trajectory, this might suggest different mechanisms of BV progression (that are driven by different bacterial species) from different healthy CSTs.

**Figure 6.**
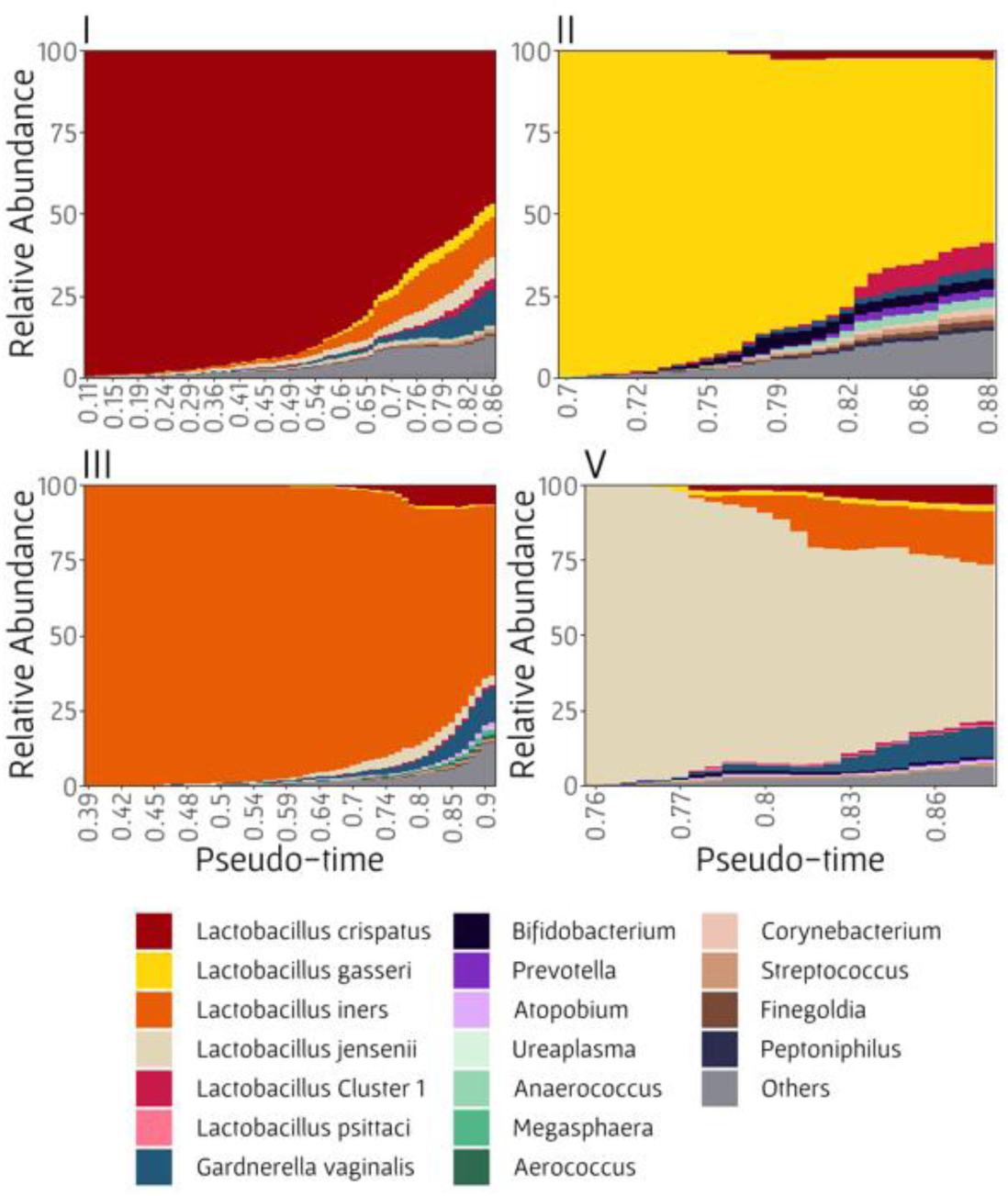
Bacterial taxa’s relative abundances as a function of by pseudo-time. Each plot represents the taxa’s abundances in each healthy manifold arm (I, II, III, and V). Taxa’s abundances were averaged using a sliding window approach to reduce noise.

## Discussion

With the goal of comprehensively characterizing the dynamics of the vaginal microbial population, we applied a framework of manifold detection inspired by common single-cell analysis. The detected manifold can serve as a map of potential routes from healthy CSTs to BV. Moreover, this approach allows us to assign each sample with a pseudo-time label, describing the location of each sample on the manifold in relation to healthy vs BV states, hence offering a proxy for the status of the sample. Our findings provide insights into the structure of vaginal microbiome dynamics and shed light on common vaginal trajectories and BV etiology, while offering an ability to evaluate the state of new samples.

Our analysis of the obtained pseudo-time labels revealed a robust association between these labels and multiple BV-related clinical parameters. With that in mind, it is worth noting that pseudo-time labels are obtained based on the bacterial composition alone, suggesting that our framework can be used to provide complementary and valuable insights into the health status of a given sample, and to infer the condition of a sample with respect to BV development stage.

Notably, amongst the BV indicators we examined, vaginal fluid exhibited only limited association with pseudo-time. Indeed, as shown in the original PVI study^29^, only BVAB1 was associated with abnormal vaginal fluid in the PVI dataset, in contrast to the PHS dataset, suggesting that this symptom needs to be further explored in different contexts. Specifically, the PVI dataset was obtained mostly from women with African ethnicity, and given the observed variation in the vaginal microbiome across ethnicities^3^, a larger dataset of more diverse ethnicities may be required. Unfortunately, the majority of samples in our study lacked accessible ethnicity data or reported ethnicity using different labeling systems, impeding our assessment of the impact of ethnicity in relation to pseudo-time.

Our analysis of microbial compositions along the vaginal microbiome manifold further revealed bacterial taxa whose abundances change in association with along pseudo-time in the different manifold arms, potentially highlight the role of various microbes in BV development. Interestingly, while the abundances of *G. vaginalis, Finegoldia,* and *Streptococcus* increased with pseudo-time in all arms, shifts in the abundances of other taxa arm-specific, suggesting different mechanisms of transition from the distinct *Lactobacillus* CSTs towards BV. *G. vaginalis* was considered associated with BV, however, its association with BV was shown to be strain-dependent^40,41^, therefore, further analysis, for example using shotgun sequencing, is required to evaluate its contribution to BV development. *Finegoldia* and *Streptococcus* were also previously identified as BV-related bacteria, together with many of the arm-specific taxa we report in our analysis^28,29,42^, yet, the relation of the arm-specific taxa to BV development from a specific CST source needs to be further evaluated. Notably, *Prevotella* – a taxon that has been hypothesized to have a dominant role in BV development (e.g., by creating a biofilm with *G*. *vaginalis*^20,43^) – appeared in our analysis only in one *Lactobacillus*-dominant arm (II), calling for further investigation into its role in BV development from other healthy CSTs. Other species which increased with pseudo-time might grow with *G. vaginalis* to form a BV biofilm, including *Finegoldia*^44^, *Streptococcus*^45^ and *Atopobium*^46^. These observations suggest that examining the progression of BV from the perspective of the previous healthy state is essential for gaining insights into the interactions between bacterial taxa and the changes in their abundance in the formation of BV biofilm.

While the observed associations between pseudo-time labels and various clinical parameters attest to the validity of our approach, it is important to acknowledge the limitations of the manifold framework. Specifically, it is worth noting that pseudo-time labels are ultimately based on changes in the bacterial composition of the microbiome, and hence correlations between pseudo-time and measures closely related to the bacterial population (such as Shannon diversity or Nugent score) are perhaps not surprising. As such, our framework and the obtained pseudo-time labels can be viewed as a powerful approach to summarize the complex and highly dimensional microbiome data, into a meaningful, clinically-relevant metric, offering a spatial perspective (i.e., manifold arm and location along the arm) for these data. Another limitation in our framework is the need to define root samples for the pseudo-time analysis. For our analyses, we used Amsel’s positive samples with high diversity as root samples, however, the ambiguous definition of BV, along with missing information in our metadata, make it difficult to determine the ideal root samples that should be used to represent BV. Additionally, even though the datasets utilized in this study provide a massive number of samples, they may still not describe the full range of variation that may be present in the vaginal human population. A larger and more diverse dataset may thus be necessary to characterize the vaginal microbiome manifold more precisely. Lastly, an important issue to consider in assessing the utility of manifold detection approaches for studying the vaginal microbiome is the marked differences between the properties of microbiome data and that of single-cell transcriptome data, to which manifold detection approaches are generally applied. Specifically, while single-cell data often describe a pre-defined, well-characterized process (such as cell differentiation), the dynamics of microbial populations are generally more noisy, including abrupt changes, exogenous perturbations, and individual-specific variation. Furthermore, microbial processes are generally not strictly regulated and not necessarily directional, and hence might exhibit back and forth transitions between different states. However, despite these differences, our study suggests that a manifold detection framework can still be applied to vaginal microbiome data and to reveal valuable insights into important microbial processes.

Combined, our analyses and findings above provide a proof of concept for the use of manifold detection methods to map vaginal bacterial samples onto a low-dimensional space, enabling the characterization of potential routes from healthy vaginal states to BV and offering a deeper understanding of vaginal microbiome dynamics. Moreover, the pseudo-time labels obtained using this approach can serve as a powerful indicator of vaginal microbiome status, and enhancing our ability to assess a woman’s health, potentially facilitating early diagnosis and treatment.

## Methods

### Datasets and data processing

Five datasets were analyzed, three cross-sectional and two longitudinal. The first cross-sectional dataset, termed VCU (Bioproject PRJNA46877), is a part of the VaHMP study^47^, which includes 3,956 samples. The data was collected from women of reproductive age, and the metadata was not published. The second dataset^3^, termed AVPVC (Bioproject PRJNA46877), is composed of 396 women of reproductive age, and includes information about Nugent score, pH, age, and ethnicity. The third cross-sectional study, termed PHS^48^ (Bioproject SRA051298), includes 242 women and information regarding Amsel’s criteria such as pH, clue cells, vaginal fluid and whiff test. The longitudinal dataset^28^, termed UAB (Bioproject PRJNA208535), included 3,700 samples and partitioned into two subsets for processing, due to differences in the sequencing platform and sequenced region. The first subset includes 1,658 samples sequenced with 454 pyrosequencing of the V1-V3 region. It was collected from 25 women of reproductive age every day for 70 days, and contains data regarding menstruation, age, ethnicity, Nugent score, pH, as well as additional important BV-related information such as BV medication, Amsel’s test results, and specific BV self-reported symptoms. The second subset includes 2,042 samples from 176 women of the V3-V4 region, sequenced via Illumina. It contains data regarding menstruation, Nugent score, and pH, however, does not include BV-related information. The second longitudinal dataset, termed PVI, includes 497 samples from 84 women (PRJNA638104)^29^. Samples were collected from four clinics, three clinics in Kenya and one in Alabama. Samples were collected once in two months, with 2-7 samples per woman. This dataset includes information about specific Amsel’s criteria state, such as pH, clue cells and discharge.

Data were acquired from NCBI as fastq files and were processed via DADA2^49^, resulting in amplicon sequence variants (ASVs) at high resolution. All datasets were filtered for more than 1,000 reads in each sample and more the 100 reads for each ASV, to exclude very rare ASVs in each dataset. The filtration step resulted 3,839 samples in the VCU dataset, 385 samples in the AVPVC dataset, 185 of the PHS dataset, 3,126 samples in the UAB dataset and 495 samples from the PVI dataset. In total, 8,541 samples were processed. Taxonomic classification was assigned to each ASV using the RDP Naïve Bayesian Classifier^50^ trained with the SILVA 16S rRNA gene database^51^. Further important species classification (*Lactobacillus*, *Gardnerella*, *Prevotella*, *Atopobium*, *Shuttleworthia* and *Sneathia*) was done with speciateIT (https://github.com/Ravel-Laboratory/speciateIT). In the second cross-sectional dataset and the subset sequenced with Illumina, ASVs assigned to *L. ultunensis* were changed to *L. crispatus*, due to their observed part in the population compared to the results. Then, samples were assigned to a specific community state type (CST) using the VALENCIA algorithm^5^.

VALENCIA implements a distance-based metric to classify each sample, based on the similarity score to the centroid of CSTs identified in a reference set. All taxonomic units were transformed to relative abundances in each sample. Rare species were removed based on their prevalence (ratio of non-zero samples to total number of samples >0.0001) and their average abundance (>0.00005) across all samples, resulting in 373 taxa in the final analysis.

### A framework for vaginal microbiome manifold detection and pseudo-time labeling

Aiming to map the potential routes of the vaginal microbiome dynamics, we applied the Partition-based graph abstraction (PAGA)^25^ algorithm, which is a common approach in the single-cell RNA-sequencing field^52^. The first step of the PAGA algorithm includes dimensionality reduction using PCA. Then, PCA values are projected onto a lower dimensional space (i.e., a manifold) using UMAP^53^ (Uniform Manifold Approximation and Projection). Finally, the samples are ordered along the manifold and a pseudo-time label is assigned to each sample using an extension of diffusion maps^54^. Notably, the diffusion-map algorithm requires a pre-defined group of cells/samples denoted as root samples, which represent the beginning of the manifold. In our analysis, we defined root samples as samples with positive Amsel’s test and high Shannon diversity (>3.5). A positive Amsel’s test is the best clinical evaluation for BV and high diversity of the microbiome is also considered as a sign for an unhealthy state^5,40^. Pseudo-time is computed by defining coordinates as dominant eigenvectors of a transition matrix that describes random walks between data points, resulting in a quantitative measure of progress through the biological process. In our analysis, the vaginal microbiome composition manifold was detected using 8,026 samples of five datasets mentioned. A total of 515 samples were held out from the manifold analysis, with 485 samples selected at random from the UAB longitudinal dataset representing 12 women, and 30 women selected at random from the AVPVC dataset with one sample per woman. These samples were subsequently utilized to evaluate the projection of new samples onto the manifold as described below. To assign held-out samples with a location on the manifold and a pseudo-time label, samples were mapped onto the manifold using a nearest-neighbor approach, using Bray-Curtis dissimilarity metric^55^ based on the microbial composition of the samples. Then, pseudo-time and UMAP values of the nearest neighbor were assigned to the held-out sample for further analyses.

### Evaluation of the association between pseudo-time labels and BV indicators

To enhance the assessment of the predictive capacity of pseudo-time labels, we focused on appraising the predictive abilities of pseudo-time in conjunction with the Nugent score, Amsel’s test result, and the specific Amsel’s criteria (i.e., pH, whiff test, occurrence of clue cells, and abnormal vaginal discharge). For this purpose, all BV indicators were categorized into predefined value options. A positive Nugent score was defined as Nugent >7, an elevated pH value was defined as pH >5.5, and a positive Amsel’s test result was defined as positive symptomatic and asymptomatic Amsel’s test outcomes. The utilization of Nugent score and pH values in previous studies has established their relevance in describing the vaginal microbiome state, particularly in relation to BV^5^. The remaining Amsel’s criteria derived from the original analysis of the PVI and PHS datasets, including identifying the presence of clue cells at a rate exceeding 20% as a positive test, and categorizing thin, gray, and homogeneous vaginal fluid as indicative of abnormality. Worth to notice that BV indicators were available in different datasets, therefore Nugent score was held from the UAB study alone, pH and Amsel’s test from UAB, PVI, and PHS, and all other Amsel’s criteria (whiff test, clue cells and vaginal fluid) were obtained from both PVI and PHS studies. To evaluate the prediction of BV indicators using pseudo-time, we employed a receiver operating characteristic (ROC) curve analysis. The ROC curve allowed us to assess the performance of pseudo-time in distinguishing between positive and negative values of each BV indicator. The area under the ROC curve (AUC) was then calculated to quantify the overall predictive accuracy of pseudo-time for BV indicator value classification. In addition, we shuffled indicator’s values 100 times to generate ROC curves and AUC calculations based on the shuffled data, to demonstrate the predictive capability of pseudo-time in comparison to a random null. This analysis provided valuable insights into the effectiveness of pseudo-time in predicting BV indicators, aiding in the assessment of its diagnostic potential for BV.

### Assessing the interaction between pseudo-time labels and menstrual cycle and biogenic amines

To investigate the association between menstruation and variations in pseudo-time, we computed a pseudo-time differentiation value for each sample, representing the difference in pseudo-time between the current sample and the earlier time-point within the same woman. These pseudo-time differentiation values were categorized into two groups: (1) samples from the previous and the next two days of menstruation period, and (2) all other samples. The difference between the pseudo-time differentiation values of these two groups was evaluated using Wilcoxon test in R.

We also evaluated association of five biogenic amines (cadaverine, putrescine, spermidine, spermine, and tyramine), obtained from another study based on samples utilized in the manifold detection framework33. These biogenic amines were measured in 79 samples of 19 women, using targeted liquid chromatography-mass spectrometry (LC-MS). Due to the uneven distribution of the samples between CSTs, we have decided to divide pseudo-time into two categories for this analysis. High pseudo-time was defined as pseudo-time >0.9. Biogenic amines levels differences between pseudo-time categories were tested using Wilcoxon test in R. Bacterial taxa abundance changes along the manifold arms Samples were categorized based on their respective manifold arm, corresponding to their CST. Subsequently, the bacterial taxa were segregated into different arms based on the location of each sample on the manifold. To ensure a smoother representation of bacterial abundance changes along the manifold, taxa abundances within each arm were averaged using a sliding window approach. Specifically, the abundances were averaged over a window size of 0.2 units of pseudo-time label, with a sliding increment of 0.01 units of pseudo-time label.

## Code availability

All code used in this manuscript has been made available at https://github.com/borenstein-lab/vaginal_microbiome_manifold.

## Acknowledgments

The authors would like to thank Sujata Srinivasan for providing the sequencing files of the PHS dataset, and to the authors of the studies included in this resource, for making their data publicly available and for responding to inquires we had during the processing of the data. We also thank former and current Borenstein lab members, and specifically Efrat Muller, Yadid Algavi, and Omri Peleg, for their helpful inputs and suggestions. This work was supported in part by National Institutes of Health (grant R01AI132441), Israel Science Foundation (Grant 2435/19 to EB) and the Edmond J. Safra Center for Bioinformatics at Tel Aviv University. The funders had no role in study design, data collection and analysis, decision to publish, or preparation of the manuscript.

## Author Contributions

MTR and EB conceived the study and wrote the manuscript. MTR obtained and processed the data and performed the analysis. All authors read and approved the final manuscript.

## References

1. Alisoltani, A. et al. Microbial function and genital inflammation in young South African women at high risk of HIV infection. Microbiome 8, 1–21 (2020).

2. Ziklo, N., Vidgen, M. E., Taing, K., Huston, W. M. & Timms, P. Dysbiosis of the vaginal microbiota and higher vaginal kynurenine/tryptophan ratio reveals an association with Chlamydia trachomatis genital infections. Front. Cell. Infect. Microbiol. 8, 1–11 (2018).

3. Ravel, J. et al. Vaginal microbiome of reproductive-age women. Proc. Natl. Acad. Sci. U. S. A. 108, 4680–4687 (2011).

4. Zhou, X. et al. Differences in the composition of vaginal microbial communities found in healthy Caucasian and black women. ISME J. 1, 121–133 (2007).

5. France, M. T., et al. VALENCIA: a nearest centroid classification method for vaginal microbial communities based on composition. Microbiome 8, 1–15 (2020).

6. Aldunate, M. et al. Antimicrobial and immune modulatory effects of lactic acid and short chain fatty acids produced by vaginal microbiota associated with eubiosis and bacterial vaginosis. Front. Physiol. 6, 1–23 (2015).

7. Fettweis, J. M. et al. The vaginal microbiome and preterm birth. Nat. Med. 25, 1012– 1021 (2019).

8. Ravel, J., Moreno, I. & Simón, C. Bacterial vaginosis and its association with infertility, endometritis, and pelvic inflammatory disease. Am. J. Obstet. Gynecol. 224, 251–257 (2021).

9. Jamieson, D. J. et al. Longitudinal analysis of bacterial vaginosis: Findings from the HIV epidemiology research study. Obstet. Gynecol. 98, 656–663 (2001).

10. Mark, S. & Phillip, E. C. Human papillomavirus and cervical cancer. Rev. Quant. Financ. Account. 8, 191–209 (1997).

11. Auriemma, R. S. et al. The Vaginal Microbiome: A Long Urogenital Colonization Throughout Woman Life. Front. Cell. Infect. Microbiol. 11, 1–11 (2021).

12. Mirmonsef, P. et al. Free glycogen in vaginal fluids is associated with Lactobacillus colonization and low vaginal pH. PLoS One 9, 26–29 (2014).

13. Gajer, P. et al. Temporal dynamics of the human vaginal microbiota-Supplementary. Sci. Transl. Med. 4, (2012).

14. Muzny, C. A. et al. An Updated Conceptual Model on the Pathogenesis of Bacterial Vaginosis. J. Infect. Dis. 220, 1399–1405 (2019).

15. Song, S. D., Acharya, K. D. & Chia, N. Daily Vaginal Microbiota Fluctuations Associated with Natural Hormonal Cycle, Contraceptives, Diet, and Exercise.

16. Rosen, E. M. et al. Is prenatal diet associated with the composition of the vaginal microbiome? 243–253 (2022) doi:10.1111/ppe.12830.

17. Gajer, P. et al. Temporal dynamics of the human vaginal microbiota. Sci. Transl. Med. 4, (2012).

18. Lugo-martinez, J., Ruiz-perez, D., Narasimhan, G. & Bar-joseph, Z. Dynamic interaction network inference from longitudinal microbiome data. 1–14 (2019).

19. Baksi, K. D., Kuntal, B. K. & Mande, S. S. ‘TIME’: A web application for obtaining Insights into Microbial Ecology using longitudinal microbiome data. Front. Microbiol. 9, 1–13 (2018).

20. Muzny, C. A. et al. Identification of Key Bacteria Involved in the Induction of Incident Bacterial Vaginosis: A Prospective Study. J. Infect. Dis. 218, 966–978 (2018).

21. Li, L. et al. Computational approach to modeling microbiome landscapes associated with chronic human disease progression. PLoS Comput. Biol. 18, 1–24 (2022).

22. Tap, J. et al. Global branches and local states of the human gut microbiome de fi ne associations with environmental and intrinsic factors. 1–11 (2023) doi:10.1038/s41467-023-38558-7.

23. Moon, K. R. et al. Manifold learning-based methods for analyzing single-cell RNA-sequencing data. Curr. Opin. Syst. Biol. 7, 36–46 (2018).

24. Cannoodt, R., Saelens, W. & Saeys, Y. Computational methods for trajectory inference from single-cell transcriptomics. Eur. J. Immunol. 46, 2496–2506 (2016).

25. Wolf, F. A. et al. PAGA: graph abstraction reconciles clustering with trajectory inference through a topology preserving map of single cells. 1–9 (2019).

26. O’Hanlon, D. E., Gajer, P., Brotman, R. M. & Ravel, J. Asymptomatic Bacterial Vaginosis Is Associated With Depletion of Mature Superficial Cells Shed From the Vaginal Epithelium. Front. Cell. Infect. Microbiol. 10, 1–10 (2020).

27. Bommana, S. et al. Metagenomic Shotgun Sequencing of Endocervical, Vaginal, and Rectal Samples among Fijian Women with and without Chlamydia trachomatis Reveals Disparate Microbial Populations and Function across Anatomic Sites: a Pilot Study.

28. Ravel, J. et al. Daily temporal dynamics of vaginal microbiota before, during and after episodes of bacterial vaginosis. Microbiome 1, 1–6 (2013).

29. Carter, K. A. et al. Associations Between Vaginal Bacteria and Bacterial Vaginosis Signs and Symptoms: A Comparative Study of Kenyan and American Women. Front. Cell. Infect. Microbiol. 12, (2022).

30. Gajer, P. et al. Temporal dynamics of the human vaginal microbiota. Sci. Transl. Med. 4, (2012).

31. Srinivasan, S. et al. Temporal variability of human vaginal bacteria and relationship with bacterial vaginosis. PLoS One 5, (2010).

32. Srinivasan, S. et al. Metabolic signatures of bacterial vaginosis. MBio 6, 1–16 (2015).

33. Borgogna, J. L. C. et al. Biogenic Amines Increase the Odds of Bacterial Vaginosis and Affect the Growth of and Lactic Acid Production by Vaginal Lactobacillus spp. Appl. Environ. Microbiol. 87, 1–16 (2021).

34. Yeoman, C. J. et al. A Multi-Omic Systems-Based Approach Reveals Metabolic Markers of Bacterial Vaginosis and Insight into the Disease. PLoS One 8, (2013).

35. Ceccarani, C. et al. Diversity of vaginal microbiome and metabolome during genital infections. Sci. Rep. 9, 1–12 (2019).

36. Muzny, C. A. & Kardas, P. A Narrative Review of Current Challenges in the Diagnosis and Management of Bacterial Vaginosis. Sex. Transm. Dis. 47, 441–446 (2020).

37. Lev-Sagie, A. et al. Vaginal microbiome transplantation in women with intractable bacterial vaginosis. Nat. Med. 25, 1500–1504 (2019).

38. Muzny, C. A., Lensing, S. Y., Aaron, K. J. & Schwebke, J. R. Incubation period and risk factors support sexual transmission of bacterial vaginosis in women who have sex with women. Sex. Transm. Infect. 95, 511–515 (2019).

39. Vaneechoutte, M. et al. Emended description of Gardnerella vaginalis and description of gardnerella leopoldii sp. Nov., gardnerella piotii sp. nov. and Gardnerella swidsinskii sp. nov., with delineation of 13 genomic species within the genus Gardnerella. Int. J. Syst. Evol. Microbiol. 69, 679–687 (2019).

40. Petrova, M. I., Reid, G., Vaneechoutte, M. & Lebeer, S. Lactobacillus iners: Friend or Foe? Trends Microbiol. 25, 182–191 (2017).

41. Shipitsyna, E., Krysanova, A., Khayrullina, G., Shalepo, K. & Savicheva, A. Quantitation of all Four Gardnerella vaginalis Clades Detects Abnormal Vaginal Microbiota Characteristic of Bacterial Vaginosis More Accurately than Putative G. vaginalis Sialidase A Gene Count. Mol. Diagn. Ther. 23, 139–147 (2019).

42. France, M., Alizadeh, M., Brown, S., Ma, B. & Ravel, J. Towards a deeper understanding of the vaginal microbiota. 7, 367–378 (2022).

43. Randis, T. M. & Ratner, A. J. Gardnerella and Prevotella: Co-conspirators in the Pathogenesis of Bacterial Vaginosis. J. Infect. Dis. 220, 1085–1088 (2019).

44. Machado, A. & Cerca, N. Influence of biofilm formation by gardnerella vaginalis and other anaerobes on bacterial vaginosis. J. Infect. Dis. 212, 1856–1861 (2015).

45. Verstraelen, H. & Swidsinski, A. The biofilm in bacterial vaginosis: Implications for epidemiology, diagnosis and treatment. Curr. Opin. Infect. Dis. 26, 86–89 (2013).

46. Hardy, L. et al. A fruitful alliance: The synergy between Atopobium vaginae and Gardnerella vaginalis in bacterial vaginosis-associated biofilm. Sex. Transm. Infect. 92, 487–491 (2016).

47. Serrano, M. G. et al. Racioethnic diversity in the dynamics of the vaginal microbiome during pregnancy. Nat. Med. 25, 1001–1011 (2019).

48. Srinivasan, S. et al. Bacterial communities in women with bacterial vaginosis: High resolution phylogenetic analyses reveal relationships of microbiota to clinical criteria. PLoS One 7, (2012).

49. Callahan, B. J. et al. DADA2: High-resolution sample inference from Illumina amplicon data. Nat. Methods 13, 581–583 (2016).

50. Wang, Q., Garrity, G. M., Tiedje, J. M. & Cole, J. R. Naïve Bayesian classifier for rapid assignment of rRNA sequences into the new bacterial taxonomy. Appl. Environ. Microbiol. 73, 5261–5267 (2007).

51. Quast, C. et al. The SILVA ribosomal RNA gene database project: Improved data processing and web-based tools. Nucleic Acids Res. 41, 590–596 (2013).

52. Saelens, W., Cannoodt, R., Todorov, H. & Saeys, Y. A comparison of single-cell trajectory inference methods. Nat. Biotechnol. 37, (2019).

53. McInnes, L., Healy, J. & Melville, J. UMAP: Uniform Manifold Approximation and Projection for Dimension Reduction. (2018).

54. Haghverdi, L., Büttner, M., Wolf, F. A., Buettner, F. & Theis, F. J. Diffusion pseudotime robustly reconstructs lineage branching. Nat. Methods 13, 845–848 (2016).

55. Beals, E. W. Bray-curtis ordination: An effective strategy for analysis of multivariate ecological data. Adv. Ecol. Res. 14, 1–55 (1984).

