## Supplementary files for "A Manifold-Based Framework for Studying the Dynamics of the Vaginal Microbiome"

### Supplementary figures

| 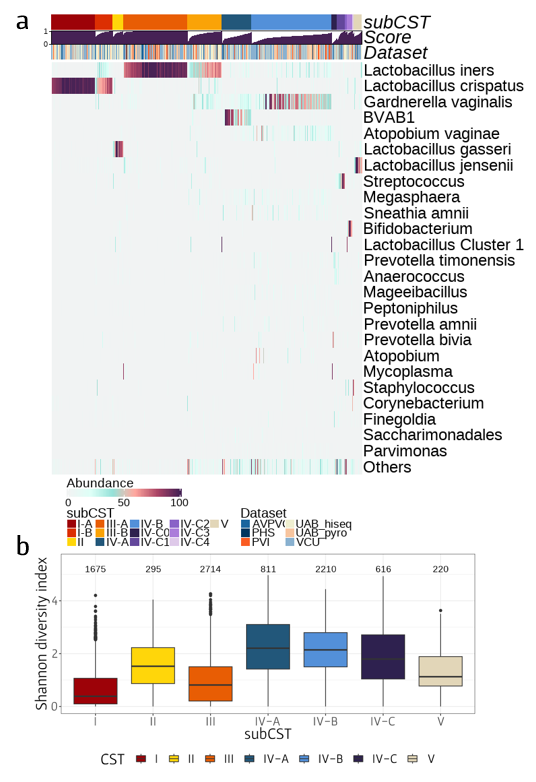 |
| --- |
| **Figure S1: (a)** **Heatmap representation of the abundances of the 25 most prevalent bacterial taxa**, where each column displays a sample and each row displays a specific bacterial taxon. The first rows of the heatmap indicates other characteristics of each sample. In the first row, each color represents the dataset the sample was obtained from. The second row's colors represent the sample's subCST assignment. In the third row, the height of the bar represents the similarity score of the samples composition to core CST defined by France et al. (see Methods). **(b)** **Box plot displays the average of Shannon diversity index for each CST**. Above each box displayed the number of samples assigned to this CST. |

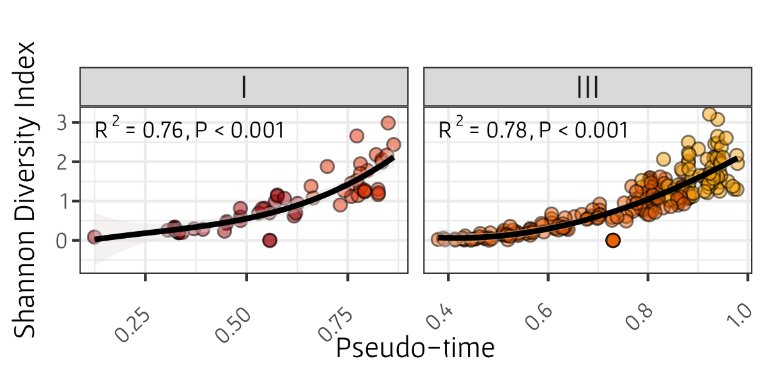

**Figure S2: Shannon diversity index as a function of pseudo-time label** of held-out samples. Pseudo-time label was assigned to each sample by its nearest neighbor on the manifold. Each plot represents a different CST arm, and colors indicate subCST assigned for each sample. The slope was determined using linear regression to transformed polynomial values of pseudo-time. The R^2^ and p-value presented in each panel were determined by Spearman correlation test. CST II and V are omitted due to each having only one held-out sample assigned.

|  | 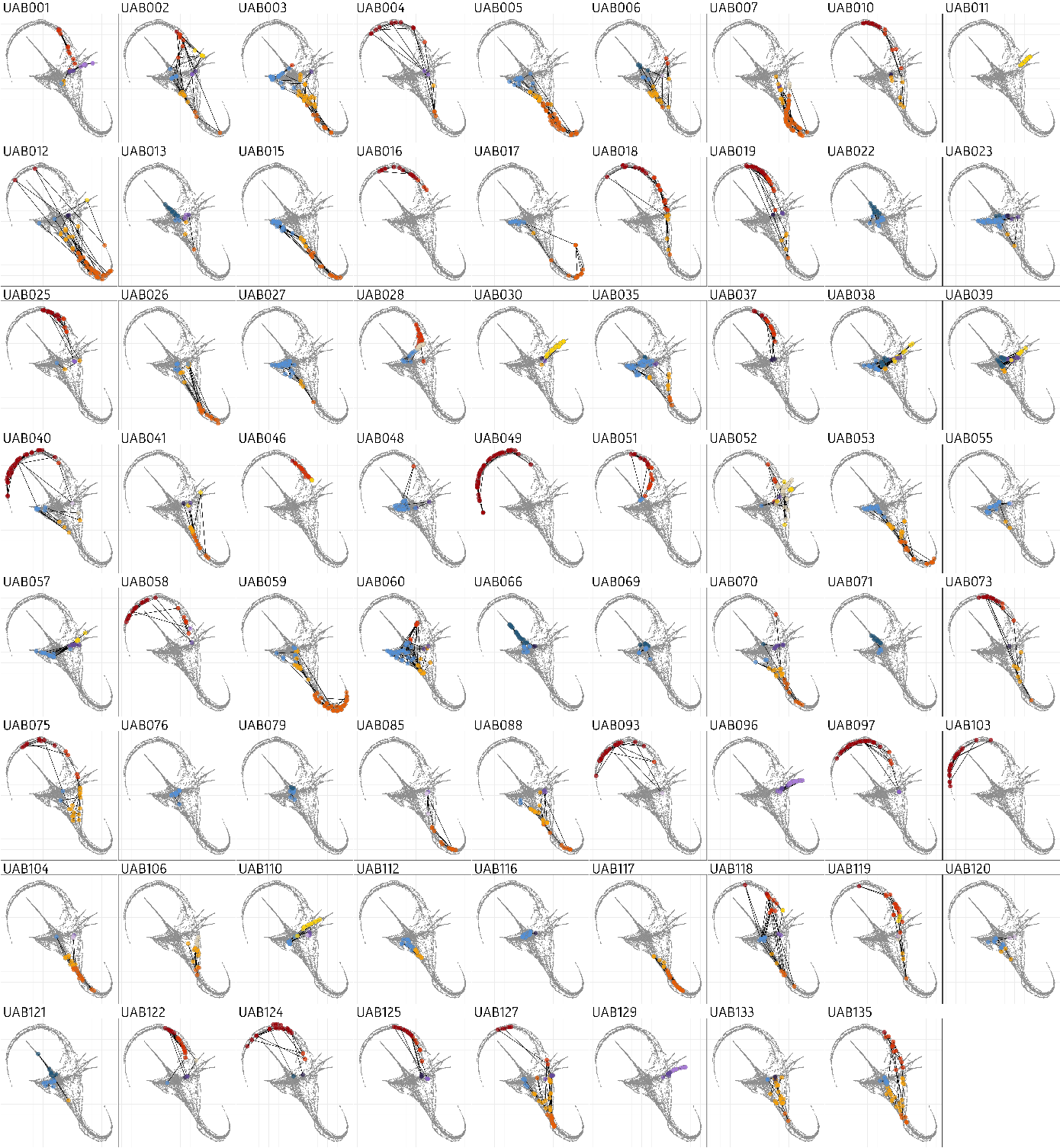 |
| --- | --- |
| **Figure S3: Trajectories of 71 women from the longitudinal dataset (UAB)** which were included in the manifold analysis. Each plot represents a different woman's trajectory. Grey dots represent the UMAP visualization of the vaginal microbiome composition manifold, and the colored dots represent the locations of samples obtained from a specific woman, where each color represent the determined subCST of the sample. Black arrows display the direction of the woman's trajectory, showing the order of the samples based on the time they were obtained in the experiment. | |

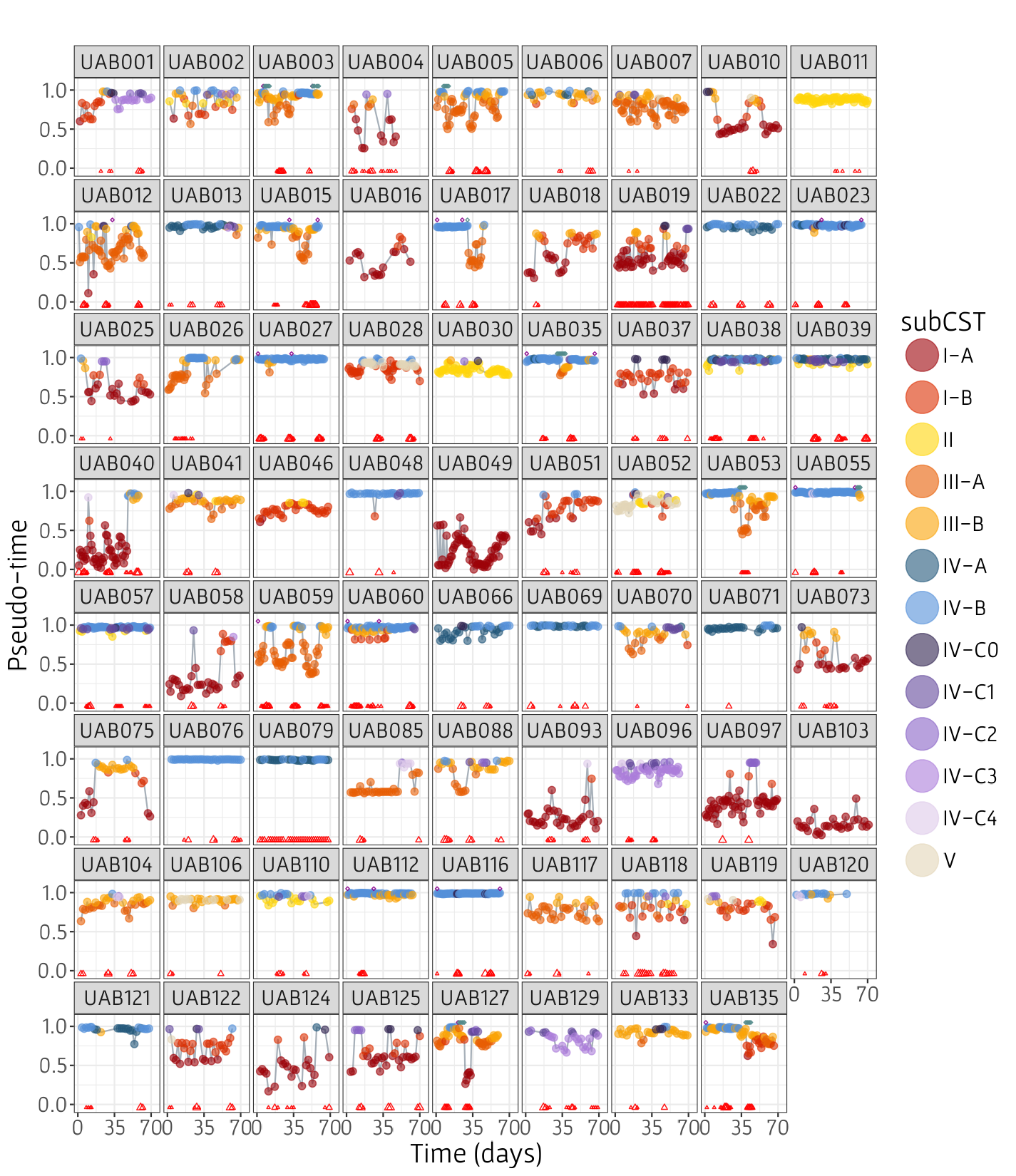

**Figure S4. Pseudo-time label to chronological time-point of each woman**, representing her trajectory. Higher value of pseudo-time label indicates it's close to root samples (i.e., BV state), while lower value of pseudo-time indicates it's distant from root samples. Dot's colors represent the sample's subCST. Red triangles in the bottom of each plot represent self-reported menstruation, where size indicates menstruation spotting (small), medium or heavy bleeding (large). Purple diamonds of the top of each plot represent time in which that sample was diagnosed with positive Amsel's criteria (with or without self-reported symptoms), and turquoise diamonds represent time with BV-medication use.

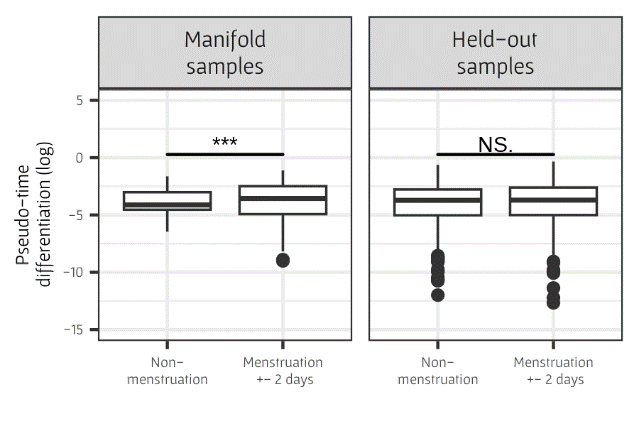

**Figure S5: Box plots of pseudo-time difference from previous sample in log scale between samples in different menstruation status**. Menstruation relates to samples from two days previous and next to menstruation period and menstruation period samples. Right panel represents results of samples from manifold analysis and right panel represents results of held-out samples. Results of Wilcoxon test are presented on top of each plot, where three asterisks indicate P. value under 0.001, two asterisks indicate P. value under 0.01, one asterisks indicates P. value under 0.05, and NS indicate non-significant.

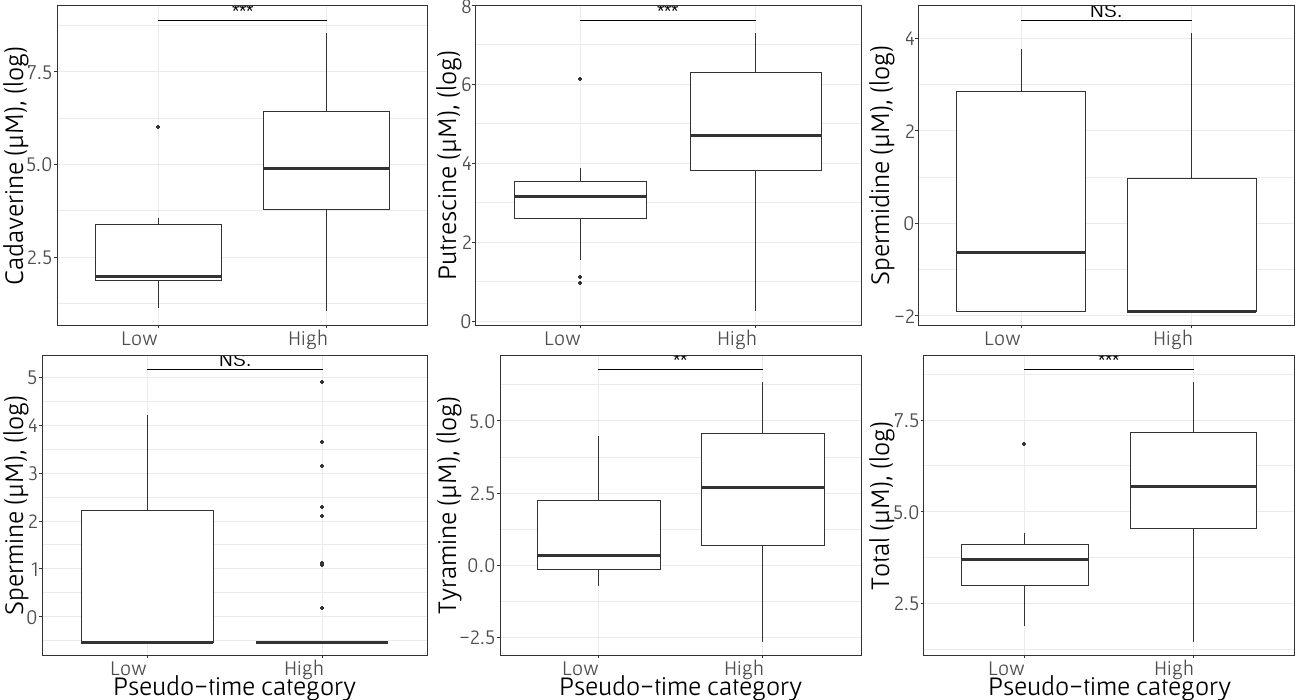

**Figure S6: Box plots of five biogenic amines levels** **in log scale**: cadaverine, putrescine, spermine, spermidine, tyramine, and total sum of all metabolites, in high pseudo-time (>0.9) and low pseudo-time. Results of Wilcoxon test are presented on top of each plot, where three asterisks indicate P. value under 0.001, two asterisks indicate P. value under 0.01, one asterisk indicates P. value under 0.05, and NS indicate non-significant.

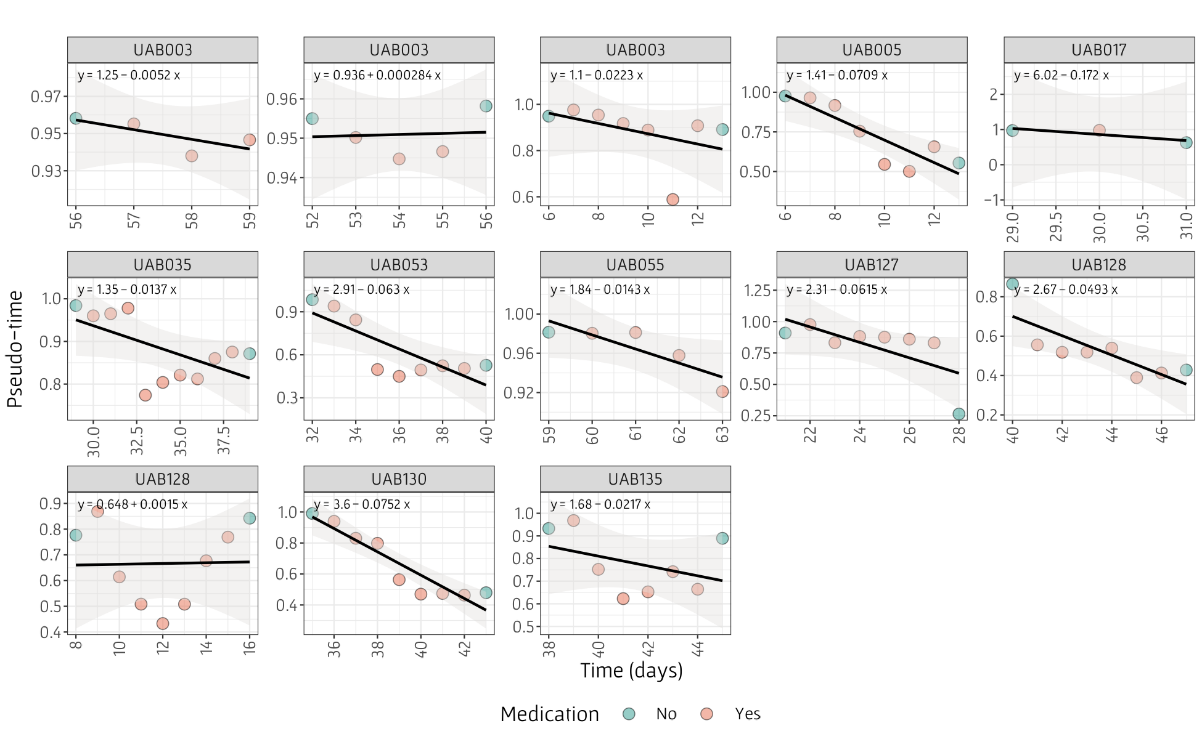

**Figure S7. Pseudo-time as a function of time in days with BV medication use**, as well as the day before and after treatment. Red points represent samples where the woman has reported medication use and green points represent the days before and after treatment. To assess the direction of pseudo-time progression, we determined the slope of each plot by employing a linear regression model equation using the `stat_poly_eq` function from the *ggpmisc* package in R. Linear model equation is presented in each top right plot, while linear regression model is represented by black lines.

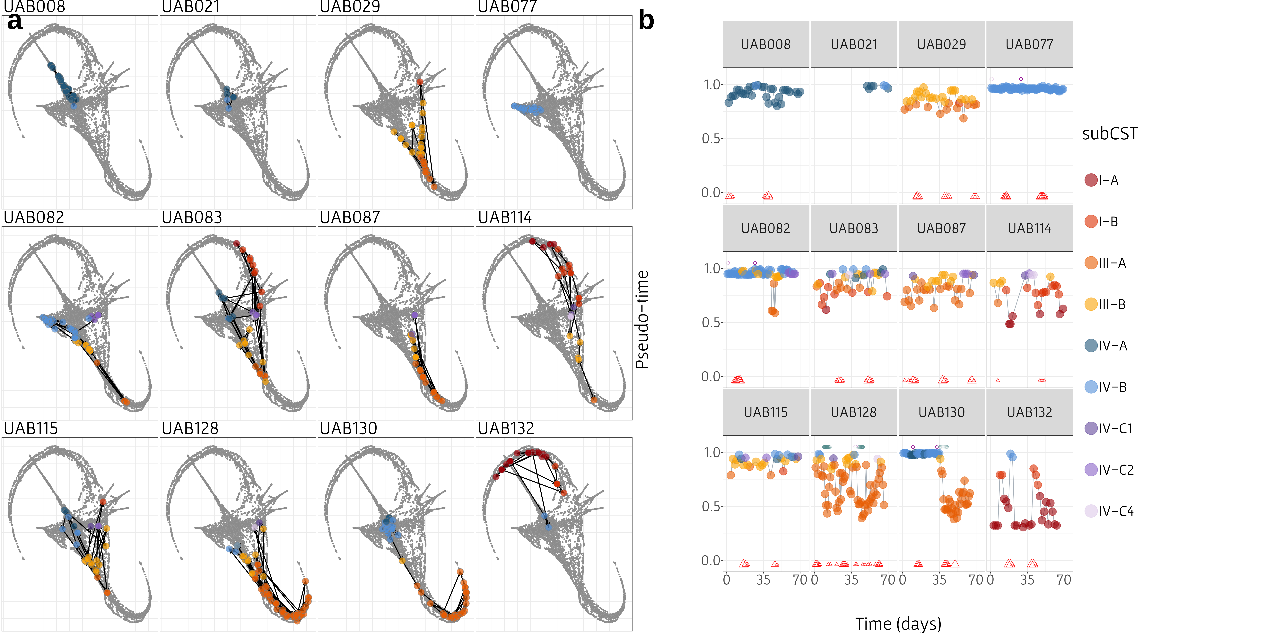

**Figure S8: Representations of the 12 women from the longitudinal dataset (UAB) which were held-out from the manifold analysis** and assigned with pseudo-time by their nearest neighbor on the manifold. **(a)** Each plot represents a different woman's trajectory. Grey dots represent the UMAP visualization of the vaginal microbiome composition manifold, and the colored dots represent the locations of samples obtained from a specific woman, where each color represent the determined subCST of the sample. Black arrows display the direction of the woman's trajectory, showing the order of the samples based on the time they were obtained in the experiment. **(b)** Pseudo-time label to chronological time-point of each woman, representing her trajectory. Higher value of pseudo-time label indicates it's close to root samples (i.e., BV state), while lower value of pseudo-time indicates it's distant from root samples. Dot's colors represent the sample's subCST. Red triangles in the bottom of each plot represent self-reported menstruation, where size indicates menstruation spotting (small), medium or heavy bleeding (large). Turquoise diamonds represent time with BV-medication use.

### Supplementary Tables

| **Supplementary Table 1: Statistical analysis results of BV indicators levels in comparison to pseudo-time value** | | | | |
| --- | --- | --- | --- | --- |
| **Parameter** | **CST** | **# of samples** | **P. value** | **FDR** |
| Nugent (Spearman) | I | 683 | 9.91E-24 | 3.47E-22 |
|  | II | 183 | 0.000896 | 0.002851 |
|  | III | 554 | 1.72E-20 | 2.01E-19 |
|  | IV-A | 246 | 1.47E-05 | 5.14E-05 |
|  | IV-B | 346 | 3.2E-06 | 1.25E-05 |
|  | IV-C | 188 | 0.031926 | 0.058811 |
|  | V | 60 | 0.010309 | 0.02255 |
| Amsel's test (Wilcoxon) | I | 209 | 0.006225 | 0.014524 |
|  | III | 747 | 1.43E-12 | 1E-11 |
|  | IV-A | 79 | 0.011699 | 0.024086 |
|  | IV-B | 771 | 1.03E-10 | 6.02E-10 |
|  | IV-C | 124 | 0.055427 | 0.08818 |
| pH (Spearman) | I | 870 | 0.053204 | 0.08818 |
|  | II | 210 | 0.51537 | 0.563686 |
|  | III | 1214 | 8.48E-23 | 1.48E-21 |
|  | IV-A | 290 | 0.20754 | 0.269034 |
|  | IV-B | 1051 | 0.005974 | 0.014524 |
|  | IV-C | 315 | 0.001564 | 0.004561 |
|  | V | 105 | 0.391245 | 0.472193 |
| Clue cells (Wilcoxon) | I | 59 | 0.075055 | 0.114214 |
|  | III | 282 | 2.75E-08 | 1.38E-07 |
|  | IV-A | 72 | 0.514457 | 0.563686 |
|  | IV-B | 236 | 0.144636 | 0.194702 |
|  | IV-C | 26 | 0.005385 | 0.014497 |
|  | V | 2 | 1 | 1 |
| Whiff test (Wilcoxon) | I | 59 | 0.030998 | 0.058811 |
|  | III | 282 | 3.53E-15 | 3.08E-14 |
|  | IV-A | 72 | 0.129594 | 0.188473 |
|  | IV-B | 236 | 0.134624 | 0.188473 |
|  | IV-C | 26 | 0.835773 | 0.886426 |
|  | V | 2 | 1 | 1 |
| Vaginal fluid (Wilcoxon) | I | 56 | 0.03807 | 0.066623 |
|  | III | 276 | 2.3E-07 | 1.01E-06 |
|  | IV-A | 71 | 0.474351 | 0.55341 |
|  | IV-B | 235 | 0.387602 | 0.472193 |
| Each BV indicator was compared to pseudo-time value in each CST arm separately. Binary BV indicators associations were calculates using Wilcoxon test, while continuous BV indicators associations were calculated using Spearman correlation. Multiple comparisons correction was conducted using FDR correction. | | | | |
